# Contextualizing Pan-Tropical Allometric Models for Biomass Estimation

**DOI:** 10.64898/2025.12.16.694295

**Authors:** Eustache Diemert, Anaëlle Dambreville

## Abstract

Allometric Models (AMs) play a central role in monitoring and mitigating climate change as they provide accurate estimation of biomass and carbon sequestered by trees from non-destructive, easy to obtain physical measurements. Unfortunately, practitioners spend considerable effort in researching, qualifying and choosing AMs for specific growth conditions. To overcome this situation Chave et al. (2014) developed a pan-tropical AM with equivalent accuracy to local, site-specific AMs.

We build upon this work to study how contextualizing AMs can improve predictive power but also provide safety checks for their application.

Our first contribution is a family of Machine Learning (ML) models that incorporate additional context pertaining to growth conditions. Evaluation shows statistically significant improvements in predictive power over a range of metrics. These models bring additional choice for practitioners in important applications such as national forest inventories, carbon certifications and calibration of satellite based biomass maps to field data.

Our second contribution proposes a principled method to estimate how much additional error one can expect when applying a given AM under new, shifting conditions - without access to ground truth biomass measurements. This method provides practitioners with a practical, data-driven safety check to qualify the risk of AMs usage in new study sites.

## 1 Introduction

AMs support numerous important applications such as National Forest Inventories (NFIs), local afforestation and reforestation projects driven by carbon markets and large-scale scientific efforts to monitor the Earth system. In typical instances AMs are used directly to convert trees measurements in the field into biomass. Recently, this has been combined with Earth Observation (EO) systems that provide Above-Ground Biomass (*AGB*) maps. These maps rely indirectly on AMs as *AGB* models predicting *AGB* from air or space-borne canopy measurements are calibrated to *AGB* estimates provided by NFIs (Hunka et al., 2024; Santoro and Cartus, 2025). As such, even small improvement in AMs estimation accuracy can lead to potentially huge corrections, as the scale of application encompasses millions of tons of biomass and sequestered carbon (Blaufelder et al.).

Tree allometry has a long history, with published works from as early as 1947 and first AMs compilations published in the 1980’s (Smith and Brand, 1983). More recently the Globallomtree database (Henry et al., 2013) registered 2,640 different equations. This research field grew as modern computing and data became more and more accessible. Nowadays, AMs are available for a variety of conditions affecting tree biomass including genetic (e.g. tree species), environmental (e.g. temperature, light, soil) and management-related (e.g. agroforestry system). Efforts have been made to consolidate them in general purpose databases (Henry et al., 2013) or more focused reviews, some of them encompassing several hundreds equations for a single country (Rojas-García et al., 2015). At the same time, the scope of application of AMs expands, leading scientists to question how to properly select an AM for specific growth conditions (Van Breugel et al., 2011), as different AMs lead sometimes to very different estimations (Zhao et al., 2012). In practice, scientists and engineers select AMs developed in the closest possible conditions (Lovell and MacKenzie, 2014), as developing new AMs incur cutting, drying and weighing at least 40 trees, a cost that few projects can afford (Návar, 2010). However, there is to the best of our knowledge no quantitative method that can evaluate if one can safely use an AM developed for another context.

At this point, we’d like to draw attention to recent developments that highlight an opportunity to improve applicability and accuracy of AMs. First, researchers have started exploring the use of ML models to replace classical power models and reported success in regional studies (Dutta Roy and Debbarma, 2024; Wongchai et al., 2022). Second, a pan-tropical AM introduced in Chave et al. (2005) and enhanced in Chave et al. (2014) has been showed to successfully replace sitespecific AMs by a single model learned on pooled data from many sites encompassing different continents, forest types and species. Their model only relies on 3 easily accessible parameters: diameter, height and wood density of trees. Considering AM development from a Statistical Learning point of view, we point out that prediction accuracy on unseen data is the result of a trade-off between data size and model complexity (Luxburg and Schoelkopf, 2008). The pan-tropical AM from Chave et al. (2014) pools data from 58 sites and learns a 3 parameter power model. Another pioneer contribution of this work is the use of *E*, an ecological pressure variable in models translating diameter to height, which can be seen as a first contextualization. Related yet not directly applicable to biomass prediction, Wang et al. (2023) showed that 8 contextual variables can help an ML model to predict coefficients of 612 AMs of the same, simple algebraic form. Following these lines of thought, we postulate that incorporating additional *contextual information* in the model would grow complexity yet has a chance to enhance prediction accuracy by better describing factors affecting biomass production. This idea is illustrated in Figure 1.

**Figure 1.**
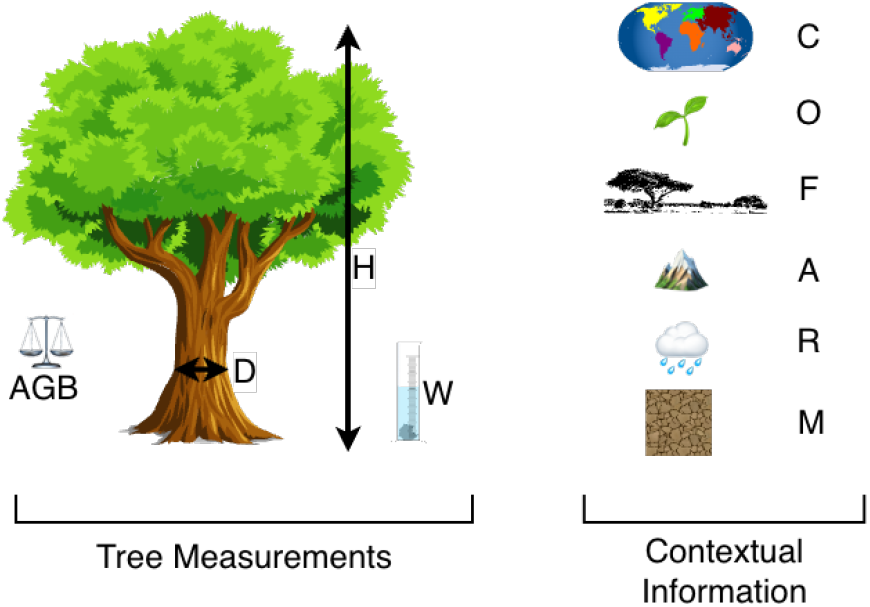
Tree measurements and contextual variables in the biomass estimation problem. Variables described in Table 1.

On the question of choice between available AMs, another insight from statistical learning is that the shift in data distributions between the calibration site (used to fit model coefficients) and application site (where inference takes place) might derail generalization of a model and produce inaccurate estimates. This situation is illustrated in Figure 2 and commonly known in statistical learning as Domain Adaptation (DA) (Cortes and Mohri, 2014). Here, we propose to use distribution distance metrics to quantify the shift (Ben-David et al., 2010; Richard et al., 2020) between calibration and application sites and learn a simple approximation of the incurred, additional biomass prediction error.

**Table 1.** Problem Variables.

| Tree Measurements |  |
| --- | --- |
| $D$ | diameter at breast height ($cm$) |
| $H$ | total height ($m$) |
| $W$ | wood density ($g.cm^{-3}$) |
| Contextual Information |  |
| $C$ | continent (*categorical*) |
| $O$ | old growth (*binary*) |
| $F$ | forest type (*categorical*) |
| $A$ | altitude ($m$) |
| $R$ | annual rainfall ($mm$) |
| $M$ | dry months (0 – 12) |
| Tree Biomass (prediction target) |  |
| $AGB$ | above ground dry matter ($kg$) |

**Figure 2.**
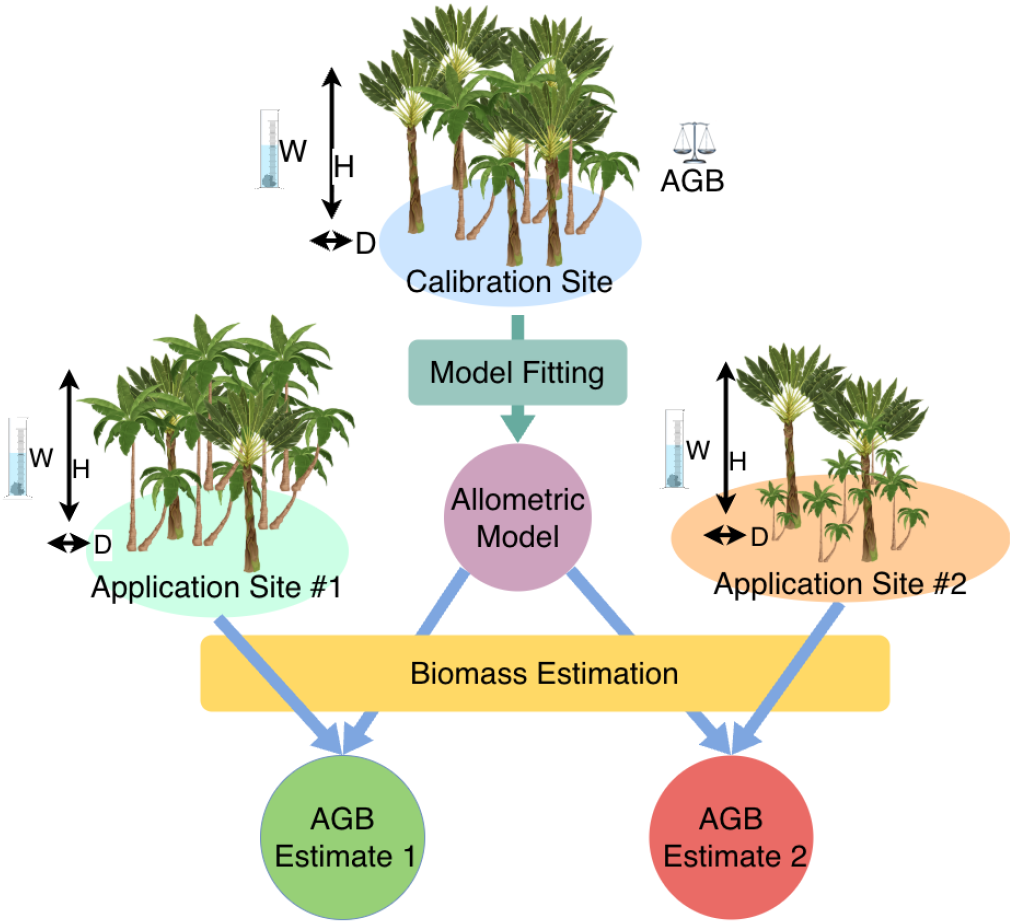
Illustration of how data distribution shift between Calibration site and two Application sites may affect estimated Biomass: a severe shift at site 2 jeopardizes estimated *AGB* accuracy while a mild shift doesn’t affect site 1 accuracy.

Our contributions can be summarized as follows:

1. 3 Contextual AMs: *ContextualChave, COFARM, COFARM-NN*, with open source code and pre-trained weights; *COFARM* improves significantly over Chave et al. (2014) on all metrics
2. *DistriCheck*, an original method to estimate additional error when using AMs on new application sites *without access to ground truth biomass of the target sites*, along with a data-driven decision procedure

## 2 Material and methods

### 2.1 Data

The dataset from Chave et al. (2014) contains measurements of trunk diameter at breast height *D* (cm), total tree height *H* (m), wood density *W* (*g*.*cm*^−3^), and total oven-dry above ground dry matter *AGB* (kg) for 4004 trees across 58 sites worldwide. We filter the dataset to focus on the portion of sites for which additional covariates are provided in Chave et al. (2005) and described in Table 1. We exclude sites that were found to contain unreliable data and later removed in Chave et al. (2014) to obtain a contextual dataset of 1,481 trees that is used for all experiments. Trees count for each site can be found in Table 2.

**Table 2.**
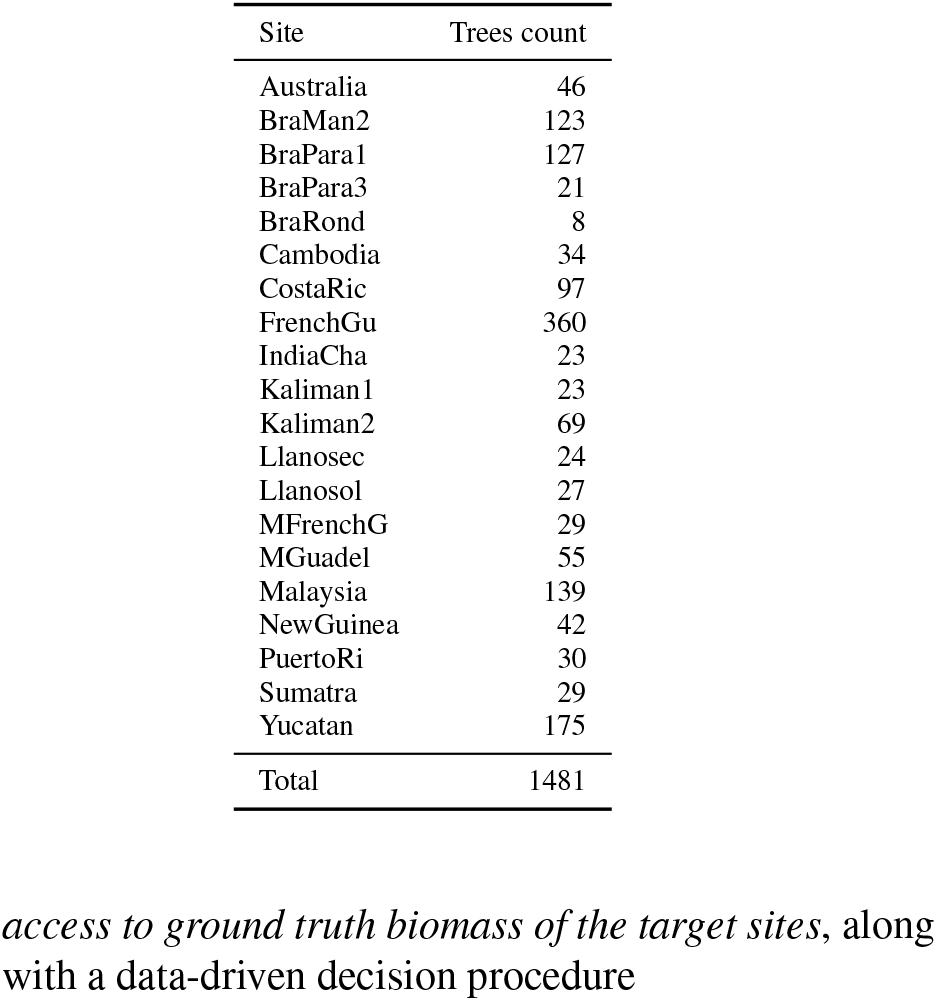
Dataset Summary.

### 2.2 Contextual AM Models

#### 2.2.1 Contextual Models Definition

We compare different models against Model 1 from Chave et al. (2014):

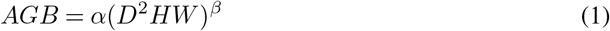

trained as explained in Eq. 2 of the original paper.

First, we select context-agnostic baselines. We train *LogReg*, a simple log-log linear regression of

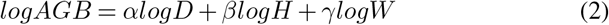

to which we adjunct an *L*^2^ regularization. We also train *HGBRT*, a Gradient Boosted Decision Trees regression model with a maximum of 1,000 iterations, access to all co-variates (tree measurements & contextual information) (Shi et al., 2022) and hyper-parameter optimization to find relevant regularization parameters.

Then, assuming contextual information can provide valuable insights to the model, we add *ContextualChave*:

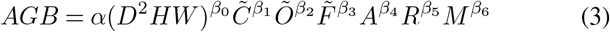

where∼designates Target Encoding (Pargent et al., 2022), a popular and efficient method to encode categorical variables.

Finally, we propose 3 deep learning models:

- *LogReg-NN*, an evolution of the *LogReg* log-log regression on *D, H, W* with 2 additional fully connected layers of 3 neurons before the final regressor
- *COFARM* a model that embeds categorical covariates *C, O, F* in 2 dimensions before pooling them with *D, H, W, A, R, M* in a log-log linear regressor
- *COFARM-NN*, inspired by wide and deep architectures (Cheng et al., 2016) pools embedded categorical co-variates *C, O, F* with *A, R, M* numerical features and passes resulting information through 3 layers of 3 fully connected neurons and ReLu blocks before the final regressor, as depicted on Figure 3.

**Figure 3.**
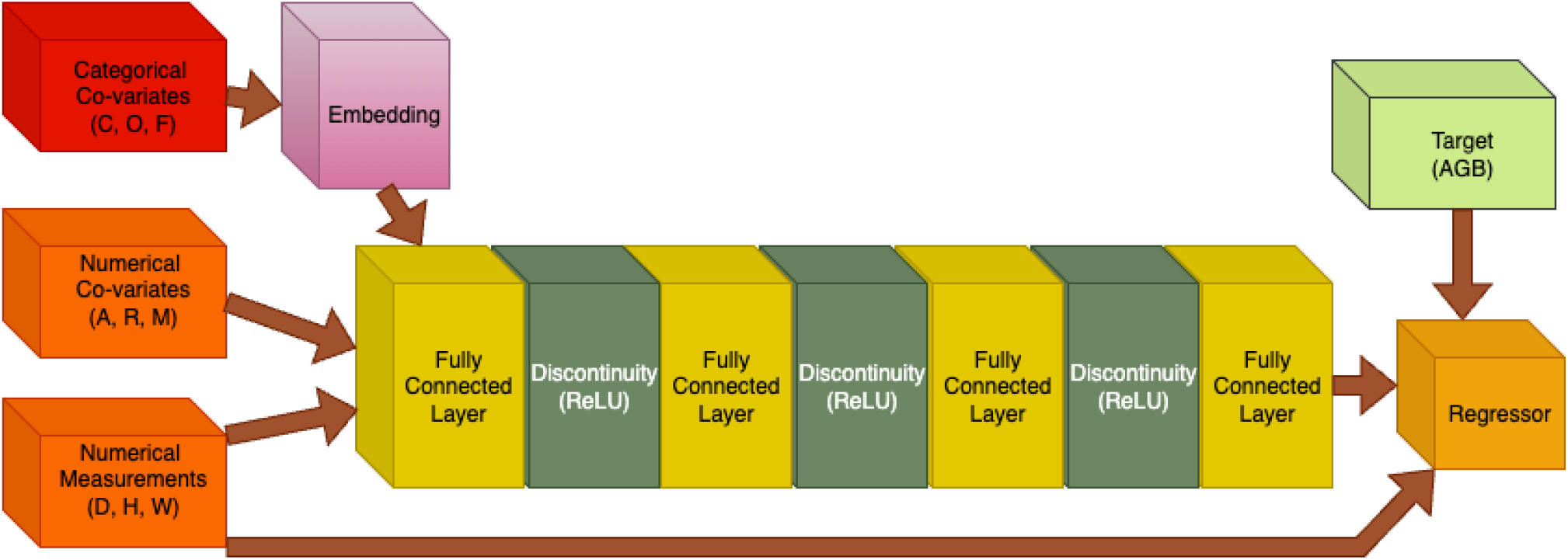
*COFARM-NN* model architecture for *AGB* prediction. Refer to Table 1 for description of variables.

These models are implemented in PyTorch (Paszke et al., 2019) and trained with the Adam optimizer (Kingma and Ba, 2017).

#### 2.2.2 Contextual Models Evaluation

First we assess *in-distribution* predictive power, simulating the case when AMs are used to estimate AGB on trees which statistical distribution coincides with the one of the calibration data. This is the primary evaluation when we assume that AMs will be used on similar (pan-tropical) sites to the ones of the Chave dataset. We thus perform 100 random train/test splits of proportion .8/.2 on the dataset of Chave et al. (2014), learning each model on train data and evaluating on hold-out test data. We report average value over the 100 splits and 90%-tile confidence intervals for each metric and model (Sileshi, 2014). Note that this methodology incorporates randomness of both data distribution and learning for non-convex methods such as deep neural networks.

Then we assess *out-of-distribution* predictive power, simulating the case when AMs are used to estimate AGB on trees which statistical distribution differs significantly from the calibration data. This concerns the case when AMs are used on sites that differ in terms of biological and environmental conditions from the ones of the Chave dataset. In this case we use a “leave one site out” protocol as described in Chave et al. (2014): we iterate over the 20 sites in the dataset and consider alternatively each one to be the test site, using the rest of the dataset as training data. Thus each iteration of the protocol contributes exactly 1 train and 1 test measurement for each metric and model. We report average value and 90%-tile confidence intervals computed over the 20 permutations for each model and metric.

Finally we define the metrics used in evaluation. For compactness we use the following notation: *y* (resp. *ŷ*) is the measured (resp. predicted) AGB, 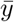 the empirical average AGB; _*i*_ and _*j*_ subscripts denote a single sample, *n* the total number of samples in the considered data partition (train or test) and *p* the number of free parameters of the model. Four metrics used in previous works are considered: Root Mean Square Error (RMSE), Residual Standard Error (RSE), Mean Absolute Error (MAE) and Coefficient of Determination (*R*^2^).

- 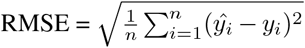
- 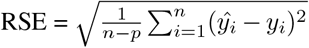
- 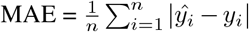
- 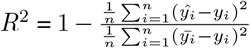

To complete the evaluation we use a Calibration Curve (Niculescu-Mizil and Caruana, 2005) that plots a moving average of the ratio between AGB predictions and observations over sorted values of AGB predictions:

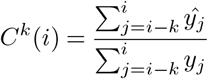

with *k* the moving window size and assuming samples are ordered by ascending AGB. This allows to observe if prediction quality varies with the prediction target, allowing to detect bias variations for smaller or bigger trees.

### 2.3 Additional Error Prediction in Shifting Conditions

#### 2.3.1 Preliminary: Unsupervised Domain Adaptation

The application of AMs to new sites can be seen as a problem of Unsupervised Domain Adaptation (DA). We observe that coefficients of an AM fitted on a calibration site may be suboptimal compared to ones that would have been fitted directly on the application site because of the difference in specific growth conditions between both sites. A key observation is that a shift in the distribution of covariates is indicative of such a situation. For instance, suppose an AM developed for a given biome and species mix in a primary forest is applied to plots from a similar biome and species mix but in a secondary forest. It is likely in this case that the AM will make a larger error in predicting *AGB* from covariates as its assumptions would be violated to a certain point when applying it on data from the application site. It is also likely that a shift in the data distribution between datasets collected at both sites will be observed, for instance larger trees could be observed in the primary forest compared to the secondary one.

We now formalize the DA problem in statistical terms following notation from Wilson and Cook (2020). We define as *source* distribution the distribution of covariates in the calibration site and *target* distribution the one of the application site. We consider a statistical model *h* : *X* → *Y*, source and target datasets *D*_*s*_ = (*X*_*s*_, *f*_*s*_) and *D*_*t*_ = (*X*_*t*_, *f*_*t*_) and a loss function *L* : ( *Y, Y* ) → ℝ. We are interested in the expected error in the target domain of a Model 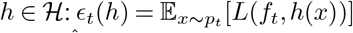. To develop a concrete model *ĥ* we assume access to an empirical source dataset 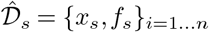 and target covariates 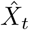; that allows to estimate the expected error of *h* on source 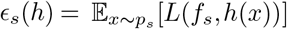 while the true target label *f*_*t*_ remains un-observed.

Many approaches have been proposed to tackle this problem, mostly in the classification case and often applied to image related tasks (Gulrajani and Lopez-Paz, 2020). For regression, we rely on Richard et al. (2020) that establish a theoretical upper bound on the expected target error *ϵ*_*t*_(*f* ) using tools from statistical learning theory. We restate their Proposition 1 for convenience:

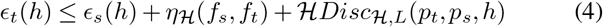

where *η*_ℋ_ (*f*_*s*_, *f*_*t*_) is the sum of the errors made by the ideal predictor on both source and target domains and *Disc*_ℋ,*L*_(*p*_*t*_, *p*_*s*_, *h*) is the hypothesis-discrepancy associated with *h* and defined as

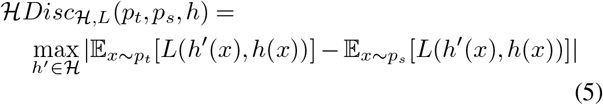

While *η* is purely related to the labeling discrepancy, ℋ*Disc* measures the discrepancy between source and target distributions, relatively to the model *h*. In other words, this bound suggests that the additional error made when applying a regression model learned on a source dataset to a target dataset can be quantified by measuring the shift between source and target covariates distributions.

#### *2*.*3*.*2 DistriCheck* : Additional Error Model

Considering that the prediction error in the target domain can be related to the shift between source and target covariates, we postulate that the suitability of an AM to new, potentially shifting conditions can be modeled. Let us consider Δ_*s,t*_*L* = *ϵ*_*t*_(*h*) − *ϵ*_*s*_(*h*), the additional error made when applying an AM to new, shifted conditions. We consider from now on *L* to be the MAE which is both telling for practical usage and respects the triangle inequality and symmetry assumptions in Richard et al. (2020). We propose a quadratic model of the additional error

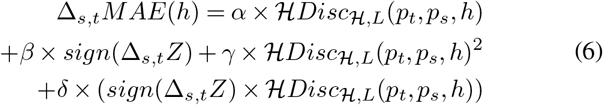

Where 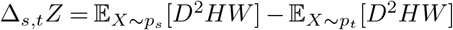. The model is learned as a Ridge regression with *L*^2^ penalty strength optimized by internal cross-validation. To estimate ℋ*Disc* in practice we solve the following constrained optimization problem

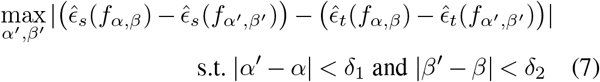

where 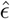 is the empirical MAE. In other words, we assume ℋ to be the family of models which parameters reside inside a ball of fixed radius centered on coefficients of the Chave model fitted on the source site data. The solution is found by solving successive quadratic optimization with the *SLS-QP* algorithm (Nocedal and Wright, 2006).

#### 2.3.3 Synthetic Distribution Shifts

A central piece of development of the additional error mode is to be able to create distribution shifts experimentally. We do so by splitting data according to contextual information to produce fake calibration and application sites. This ensures existence of a shift in tree measurement and biomass distributions between the two sub-parts of the data.

For each numerical variable (*A, R, M* ), we compute the 25^*th*^ and 75^*th*^ percentiles and use them as thresholds to divide the distribution into two sub-distributions, corresponding to source and target datasets. Table 3 shows the thresholds used to implement the shifts. For example, for altitude, the 25^*th*^ percentile is 42.0 m (“altitude-low” shift). The source dataset is thus composed of trees located < 42 m, and the target dataset encompasses the trees located ≥ 42 m. For the “drymonths-many” shift, the 85^*th*^ percentile has been used to create the shift because the 75^*th*^ percentile was creating a shift identical to the “foresttype-dry” shift. *C* (Asia or Americas) and *O* (0 or 1) have only two categories. Source and target datasets are thus created by separating the trees based on one of the two values. *F* encompasses four categories. Each one of them has been used to create a supplementary shift in distributions. We also consider two random shifts by randomly separating the dataset, resulting to a total of 14 shift types. Finally, we also implement reversed shifts by switching source and target datasets leading to a total of 28 shifts. For each of them, the objective is to train the AM from Chave et al. (2014) on the source dataset and to predict *AGB* and the additional error on the target dataset without observing target *AGB*.

**Table 3.** Thresholds used to experimentally create the shifts with Wasserstein distance and ℋ*Disc* between source/calibration and target/application.

| Var. | Shift | Threshold | Wass. Dist | $\mathcal{H}Disc$ |
| --- | --- | --- | --- | --- |
| A | altitude-high | 100.0 | 8954.7 | 1070.9 |
| A | altitude-low | 42.0 | 9736.9 | 2198.8 |
| C | continent-asia | Asia | 7019.4 | 1611.0 |
| M | drymonths-few | 1.0 | 25083.8 | 7225.7 |
| M | drymonths-many | 5.0 | 11649.6 | 12092.9 |
| F | foresttype-dry | Dry | 11524.2 | 1718.7 |
| F | foresttype-mangrove | MoistMangrove | 10632.2 | 6656.6 |
| F | foresttype-moist | Moist | 3986.4 | 489.1 |
| F | foresttype-wet | Wet | 18408.5 | 3090.1 |
| O | oldgrowth-0 | 0 | 11702.0 | 4628.4 |
| R | rainfall-high | 3125.0 | 16162.3 | 2824.7 |
| R | rainfall-low | 1862.0 | 11816.2 | 3965.9 |
| - | random-1 | - | 2400.3 | 126.8 |
| - | random-2 | - | 3954.3 | 1488.6 |

To illustrate the magnitude of the discrepancy between source and target datasets for each shift, we compute the Wasserstein distance on *D*^2^*HW* (Panaretos and Zemel (2019)). Moreover, a significant shift in distributions implies that it would be possible to predict the origin of a tree (source or target distribution) based on its covariates values (*D*^2^, *H*, and *W* ). To illustrate this statement, we train a Gradient Boosted Decision Trees classifier model (Friedman, 2002) on the Chave dataset to predict whether a tree belongs to the source dataset or not, for each shift ((Lopez-Paz and Oquab, 2018)). If it performs better than random it confirms that a shift exists and is significant.

#### 2.3.4 Additional Error Model Training and Evaluation

In order to evaluate the proposed model we first randomly split the 1481 trees of our data into *D*_*train*_ (741 trees) and *D*_*test*_ (740 trees). *D*_*train*_ is used to fit coefficients of Equation 8 while *D*_*test*_ is used to evaluate the capacity of the model to correctly predict additional error.

For the training, we randomly split *D*_*train*_ into *i* = 1 … *n* sub-datasets 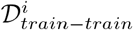 and 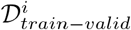 . In turn, each of these *n* sub-datasets generates *s* = 1 … *S* synthetic shifts by applying the procedure detailed in Section 2.3.3. A Chave model is then fitted on each 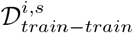 instance and inference performed on corresponding 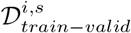 . Every (*i, s*) pair thus contributes one training sample of the form (Δ*MAE*^*i,s*^, ℋ*Disc*^*i,s*^, *sign*(Δ*Z*)^*i,s*^). With *n* = 2000 repeats and 14 shifts in 2 directions this procedure generates a dataset 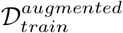 of 56000 samples. The model is then trained on a standardized version of this dataset with a square loss and *L*^2^ penalty strength optimized through cross-validation.

For the evaluation, *D*_*test*_ is first divided into 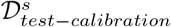 and 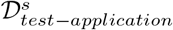 datasets based on the same *s* = 1 … *S* shift definitions. A Chave model is trained on 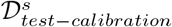 and Δ*MAE*^*s*^ computed from 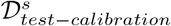 and 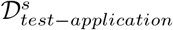 . This value is then compared to the additional error predicted by Equation 8 based on ℋ*Disc* and *sign*(Δ*Z*).

## 3 Results

### 3.1 Contextual Models Performance

We focus first on *in-distribution* performance as depicted in Table 4 and illustrated on Figure 4 for MAE. In the train partition we observe as expected a reduction of errors magnitude with more complex models. Except for *HGBRT* this trend exists as well in the test partition, indicating more complex models can generalize. A first insight is that models using additional contextual information out-perform similar size models on every test metric - except Test RSE for *COFARM-NN*, which penalizes model size. Comparing *LogReg* and *LogReg-NN* versus *ContextualChave* on test indicates a 4% decrease in MAE due only to contextual information, as models are of comparable size. A second insight is that additional model complexity doesn’t necessarily incur more predictive power without contextual information. See the performance of *ContextualChave* which can achieve a better test MAE with less parameters (164.4, *p* = 8) than *LogReg-NN* (171.4, *p* = 16). When comparing *ContextualChave* to *COFARM* we observe a 8% decrease in test MAE (-11% in RMSE), indicating approximately the uplift we can attribute to a better suited ML model using the same, contextual information. We note that this amounts approximately to double the uplift of just adding contextual information to more traditional models. Third, The *COFARM* and *COFARM-NN* family dominate on all test metrics except RSE. *COFARM* outperforms significantly Chave et al. (2014) *on every single metric* by a comfortable margin: -15% MSE, -11% RSE, -15% MAE and +4% *R*^2^. The more complex *COFARM-NN* is a close runner-up with a -14% reduction in MAE but probably suffers from its complexity (high *p*) with respect to available data size. Besides, *COFARM* produces lower bias over the whole AGB distribution as can be observed on Figure 5. Indeed, the *COFARM* calibration curve is closer to the ideal value of 1.0, especially so for small but also large *AGB* (> exp(6) = 400 kgs), an area where the original AM is suboptimal. All in all, these results provide evidence of the generalization power of the *COFARM* and *COFARM-NN* models.

**Table 4.** *In-Distribution* Predictive Power of different AMs assessed with 100 Random Train/Test Splits (bold = best, underline = second best, ^∗^ = statistical significance at 90% vs Chave et al. (2014))

| Dataset | Metric | Chave et al. (2014) | ContextualChave | HGBRT | LogReg | LogReg-NN | COFARM | COFARM-NN |
| --- | --- | --- | --- | --- | --- | --- | --- | --- |
| - | parameters | 2 | 8 | 156,000 | 4 | 16 | 28 | 132 |
| - | contextual | no | yes | yes | no | no | yes | yes |
| train | RMSE | 834.1 ± 10.5 | 828.0 ± 10.3 | <u>610.3</u> ± 16.2 * | 767.1 ± 8.4 * | 767.0 ± 8.4 * | 700.9 ± 8.9 * | **600.6** ± 19.6 * |
| train | RSE | 836.9 ± 10.5 | 839.4 ± 10.4 | - | <u>772.3</u> ± 8.5 * | 788.6 ± 8.6 * | **736.6** ± 9.4 * | 806.7 ± 26.3 |
| train | MAE | 175.4 ± 1.4 | 164.2 ± 1.3 * | **82.2** ± 1.3 * | 169.4 ± 1.3 * | 169.4 ± 1.3 * | 148.9 ± 1.2 * | <u>132.1</u> ± 2.2 * |
| train | R2 | 0.884 ± 0.003 | 0.886 ± 0.003 | <u>0.938</u> ± 0.003 * | 0.902 ± 0.003 * | 0.902 ± 0.003 * | 0.918 ± 0.003 * | **0.939** ± 0.004 * |
| test | RMSE | 792.1 ± 38.8 | 781.7 ± 41.5 | 857.2 ± 70.3 | 738.7 ± 33.2 | 738.7 ± 33.2 | **673.3** ± 34.8 * [-15%] | <u>735.0</u> ± 55.7 |
| test | RSE | 794.8 ± 39.0 | 792.5 ± 42.1 | - | <u>743.7</u> ± 33.4 | 759.5 ± 34.1 | **707.6** ± 39.6 * [-11%] | 987.2 ± 74.8 |
| test | MAE | 176.8 ± 5.4 | 164.4 ± 5.1 * [-7%] | 174.7 ± 7.6 | 171.4 ± 5.1 | 171.4 ± 5.1 | **151.0** ± 4.6 * [-15%] | <u>152.2</u> ± 5.6 * [-14%] |
| test | R2 | 0.873 ± 0.012 | 0.877 ± 0.013 | 0.863 ± 0.014 | 0.888 ± 0.010 | 0.888 ± 0.010 | **0.904** ± 0.011 * [+4%] | <u>0.890</u> ± 0.016 |

**Figure 4.**
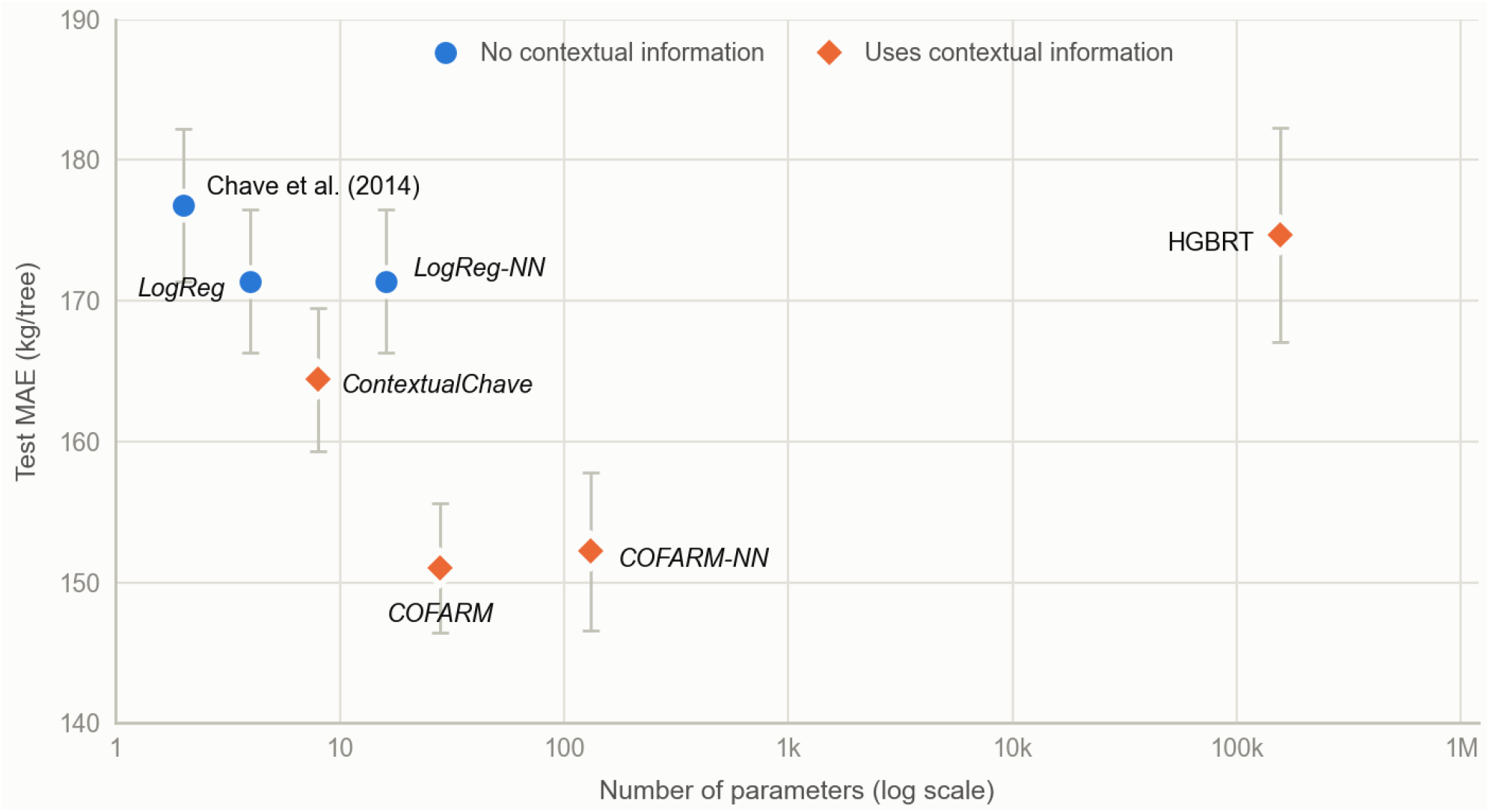
*In-Distribution* Predictive Power vs Model Complexity as measured by mean Test MAE and 90% confidence interval over 100 random train/test splits and *p*. Diamond markers indicate access to contextual variables.

**Figure 5.**
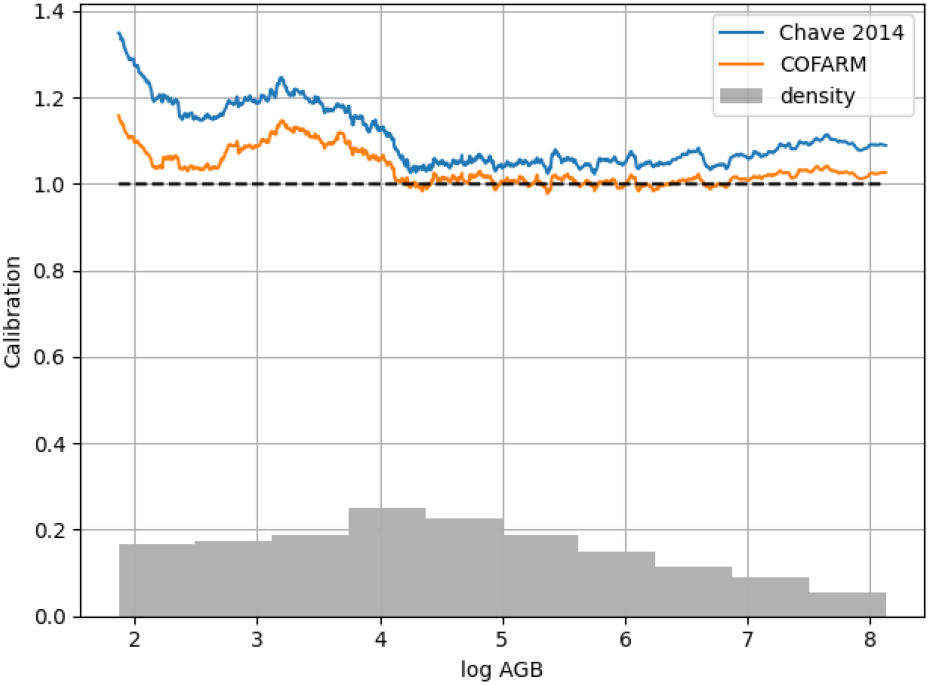
Calibration plot of *AGB* predictions for the Chave and *COFARM* models. Density refers to probability of a random sample in the Chave dataset having a specific AGB value.

A second axis of performance is generalization under shifting conditions, as measured using the Leave-one-site-out protocol. This protocol simulates *out-of-distribution*, a much more challenging case. Results in Table 5 suffer from high uncertainty since only 20 shifts can be simulated with available data. *COFARM* and *COFARM-NN* present tendency evidence of out-performing Chave et al. (2014), yet the confidence interval is so large that one cannot comfortably conclude. We note that Chave et al. (2014) didn’t compute confidence bounds with this protocol but rather use it to illustrate so-called outlier sites for which *out-of-distribution* generalization is difficult (see Fig 3 of their paper).

**Table 5.** *Out-of-Distribution* Predictive Power of different AMs assessed with 20 Leave-One-Site-Out splits (no highlighting of results as metrics variance in test partition prevents from drawing robust conclusions)

| Dataset | Metric | Chave et al. (2014) | ContextualChave | HGBRT | LogReg | LogReg-NN | COFARM | COFARM-NN |
| --- | --- | --- | --- | --- | --- | --- | --- | --- |
| train | RMSE | 831.4 ± 24.7 | 824.9 ± 20.6 | 568.8 ± 21.1 | 766.7 ± 19.7 | 769.4 ± 20.7 | 718.4 ± 17.4 | 663.0 ± 28.9 |
| train | MAE | 175.5 ± 5.1 | 164.1 ± 4.0 | 77.5 ± 1.9 | 169.7 ± 5.3 | 170.5 ± 5.7 | 152.1 ± 3.7 | 139.3 ± 3.4 |
| train | R2 | 0.884 ± 0.005 | 0.886 ± 0.004 | 0.946 ± 0.004 | 0.901 ± 0.004 | 0.901 ± 0.004 | 0.913 ± 0.004 | 0.925 ± 0.009 |
| test | RMSE | 592.7 ± 416.8 | 600.2 ± 335.9 | 849.0 ± 574.8 | 543.0 ± 354.3 | 544.1 ± 354.2 | 537.0 ± 321.4 | 631.6 ± 358.1 |
| test | MAE | 242.8 ± 177.6 | 219.6 ± 131.9 | 302.4 ± 209.2 | 224.9 ± 154.2 | 225.8 ± 154.1 | 219.2 ± 134.1 | 228.0 ± 158.6 |
| test | R2 | 0.835 ± 0.062 | 0.825 ± 0.079 | 0.705 ± 0.191 | 0.844 ± 0.065 | 0.845 ± 0.066 | 0.801 ± 0.094 | 0.661 ± 0.203 |

### 3.2 Evidence of Covariate Shift

Table 3 shows the Wasserstein distance for each shift type that has been implemented. The distance is the lowest for random shifts, meaning that there is little difference between source and target distributions for those shifts. *foresttype* − *moist* shift shows also a low distance, meaning there is only a minor shift of distribution. *Drymonths* − *few* has the highest distance, highlighting a very different distribution for the trees located in places where there is no dry months. ℋ*Disc* is also the highest for *drymonths* − *few* showing the maximal discrepancy for this shift. However, there is no linear relationship between the Wasserstein distance and ℋ*Disc*, i.e. a higher distance is not always related to a higher ℋ*Disc*. This is probably because the distance is model-agnostic and rely only on covariates distributions while ℋ*Disc* is conditional to the fitted Chave model.

For all shifts except the random ones, a Gradient Boosting Classifier is able to learn to predict when samples come from source or target datasets (data not shown, p-value < .0001). This highlights a clear shift in the covariates distributions between the source and target distributions even when the Wasserstein distance is low.

### 3.3 Generalization Error Prediction

Figure 6 shows the predicted additional MAE due to distributions shift as a function of the observed one. Our model was able to predict Δ_*s,t*_*MAE* with a high precision (R^2^ = 0.842) meaning that it was able to learn the additional error based only on ℋ*Disc* and *sign*(Δ_*s,t*_*Z*). The red area highlights where predictions would have been falsely optimistic (positive Δ_*s,t*_*MAE* predicted as negative) and potentially lead to qualifying an AM when it should not. We observe only 1 data point in this case. These results show that the *DistriCheck* model is conservative and able to predict the sign and magnitude of the additional error correctly. More details about *MAE* values for each shift can be found in Appendix B1.

**Figure 6.**
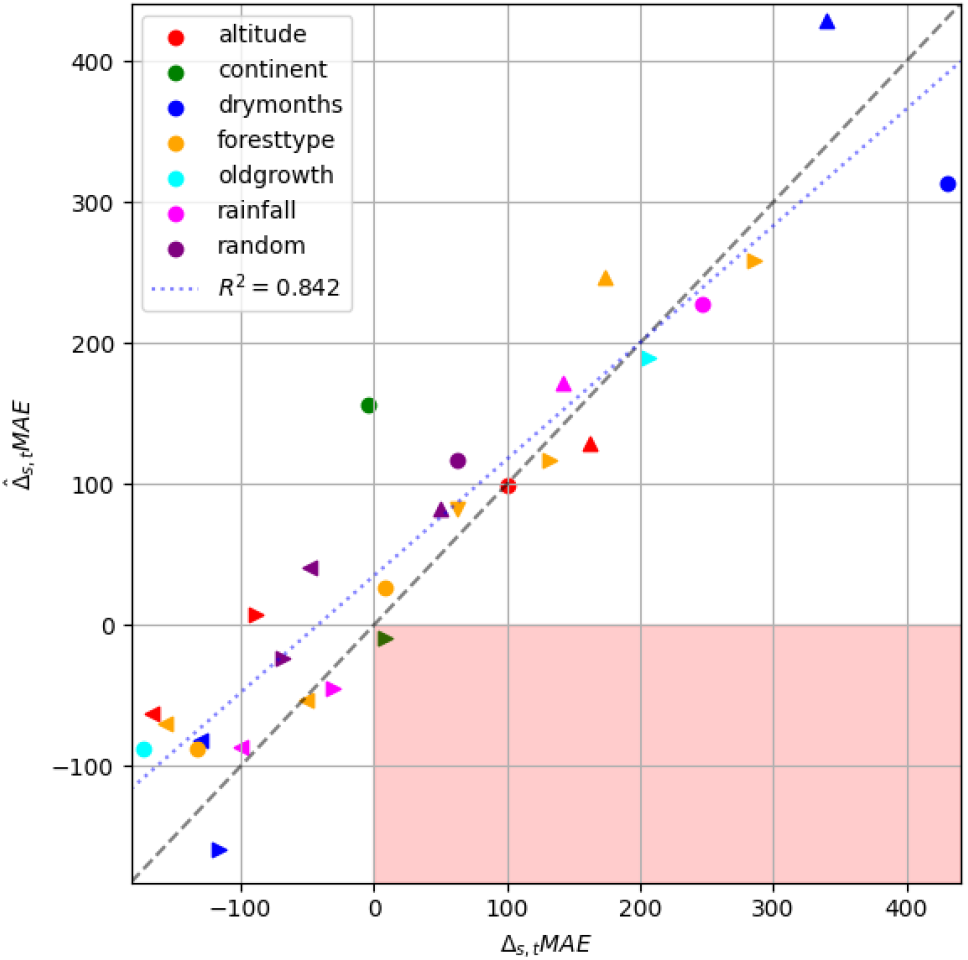
*DistriCheck* Predicted vs Actual Additional Error due to Covariate Shift. Color indicates the variable chosen to simulate the shift while marker shapes indicate different shifts magnitudes and directions. The red area highlights cases where the model predicts that the AM estimation will be more precise on the shifted data whereas the actual additional error is positive.

The final model equation stands as follows:

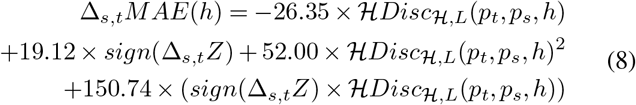

The coefficient for ℋ*Disc* is negative, however the one for ℋ*Disc*^2^ is twice as big and positive leading to a global positive effect of ℋ*Disc* on Δ_*s,t*_*MAE*. This means that Δ_*s,t*_*MAE* will get bigger when ℋ*Disc* increases. The coefficient for *sign*(Δ_*s,t*_*Z*) being positive, it means that Δ_*s,t*_*MAE* will be higher when 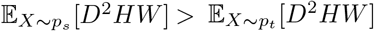, i.e. when source covariates are bigger than target covariates. The interaction term between *sign*(Δ_*s,t*_*Z*) and ℋ*Disc* shows the biggest coefficient high-lighting that when 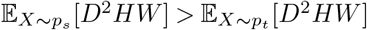 a high ℋ*Disc* will increase the Δ_*s,t*_*MAE*. On the contrary, when 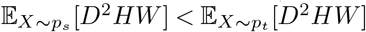 a high ℋ*Disc* will decrease Δ_*s,t*_*MAE*.

## 4 Discussion

### 4.1 Summary of Key Findings

We present a trilogy of novel, contextual models to predict *AGB* from easy to obtain tree measurements and contextual information. Among contextual models, *COFARM* improves significantly out-of-sample estimation across all metrics, therefore reducing errors magnitude by 15% compared to the established pan-tropical baseline. We show that this improvement comes partly from contextual information but most importantly from more sophisticated modeling that is able to make the most of this additional information. The proposed model is also easily amenable to incorporating additional information related to growth conditions.

Moreover, we propose *DistriCheck*, a principled method to predict the additional error incurred by changing growth conditions between the site where a given AM is learned and the one where it is applied. This method can be used with any AM given that one has access to tree measurement data from both sites (*AGB* data from target site is not required), allowing practitioners to test applicability and judge it from observed data.

### 4.2 Limitations

Limitations of present work include data, modeling and methodological considerations. First, we’d like to praise the authors of Chave et al. (2014) for the constitution of this incredibly helpful dataset. Yet, the datasets we use are still imperfect and may potentially misrepresent characteristics of tropical tree growth as they can be found in the wide area of tropical forests. In that respect, we inherit the limitations of the original work, including the applicable range of *D* (5-156cm) and the exclusion of palms and liana species. More diverse open source biomass datasets would be required to develop more general pan-tropical AMs. Experiments on adding contextual information to AMs could be conducted only on the portion of the dataset that contains such information, totaling 1,481 trees. Second, even though we took care to evaluate a significant number of ML models, including linear, tree-based and deep neural networks, we possibly missed more accurate models. Finally, our proposed error prediction method is based on established statistical learning theory, yet the generalization bounds tightness can be questioned. Even if our experiments suggest a strong link and predictability in practice, only a larger scale use of the technique would give necessary feedback. Additional studies on other potential co-variate shifts between conditions for learning and applying an AM would be of great value to the community.

### 4.3 Recommendation on Applicability of Allometric Models

In light of the experiments in Section 3.3, we propose a data-driven sanity check to decide if a given AM is suitable to be applied to a new dataset of tree measurements. The process is depicted in Figure 7 and starts with the computation of statistics on both the calibration dataset used to fit the AM and the newly considered dataset. It follows by injection of these statistics in the *DistriCheck* model (Equation 8) an estimation of the error incurred by the use of this AM in the new application site. The principal can then declare if the forecasted error falls within her pre-defined acceptable error range. If the answer is positive, the biomass predictions of the AM are judged satisfying and can be used. Conversely, an alternative AM should be considered; and if unfortunately none is available a case made to develop a new AM using destructive measurements. The advantage of this process are three-fold. First, it bases AM selection on objective, data-driven criteria. Second, it provides trust and visibility as shift statistics may be openly communicated to stakeholders. Finally, it motivates creation of new AM and the incurred costs on a strictly needed basis. We note that the proposed sanity check should be understood as a necessary yet not sufficient condition and by no means qualifies a given AM by itself, but rather adds an objective criteria of applicability. Indeed, the absence of a distribution shift on tree measurements doesn’t guarantee that the AM captured the relation of the former with *AGB* and we still recommend in addition to follow the prescriptions in each AM reference publication.

**Figure 7.**
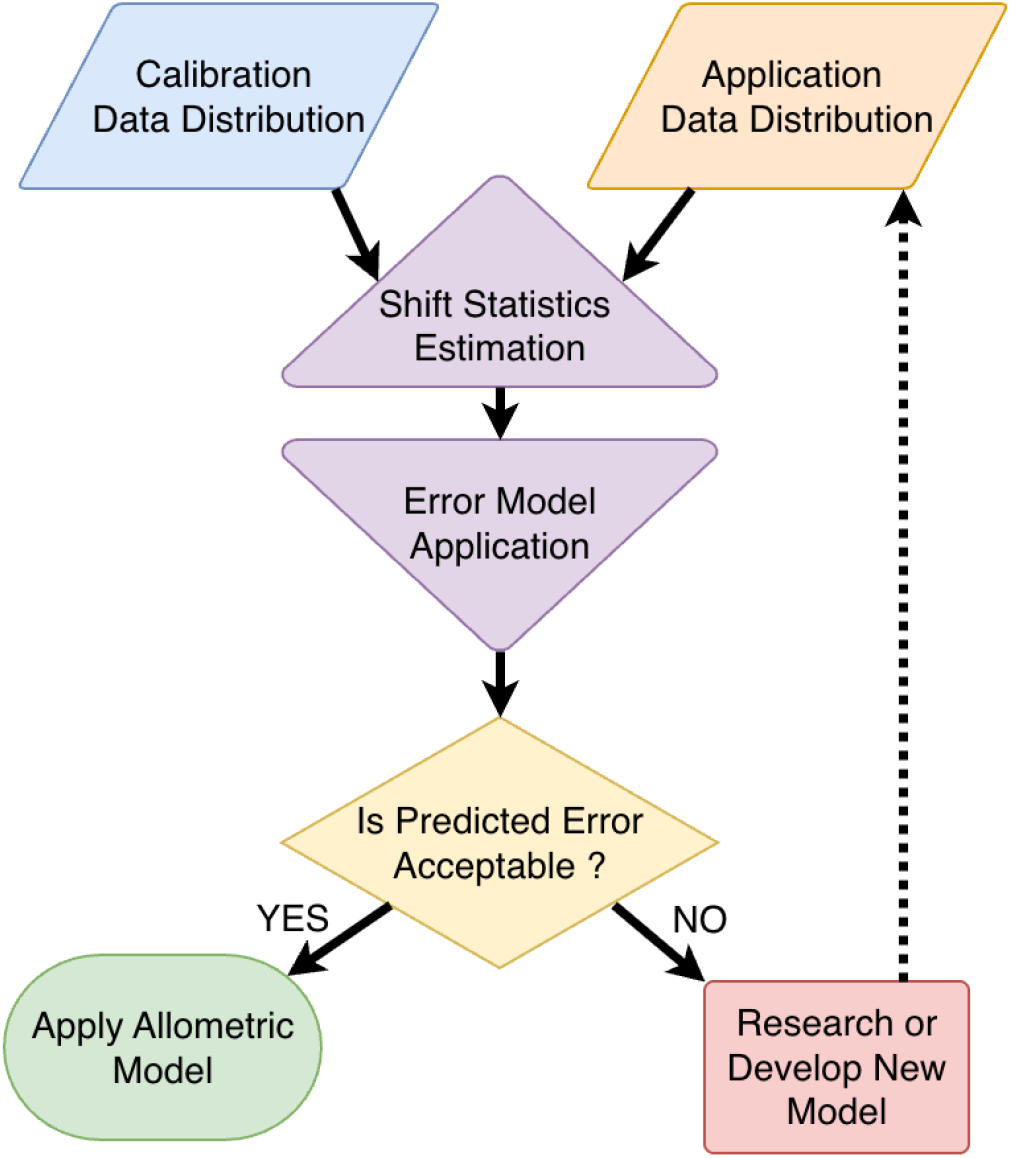
Recommended Process for Qualifying or Adjusting Allometric Models based on *DistriCheck* Predicted Additional Error.

### 4.4 Perspectives

We foresee exciting future works in many directions. On the allometric modeling side, we see recent developments in Geospatial Foundation Models (Brown et al., 2025) as an opportunity to bring even more contextual information to AMs in an efficient manner. We also anticipate that deep embeddings used to represent contextual information could well incorporate species information when it is available. On the methodological side, we wish for more raw data pertaining to AM development to be released openly. Not only would it allow more modeling research to happen, but also could increase reliability of AM choices by generalizing the kind of safety checks we propose to all AMs. More importantly, as a scientific field, open code and data is a formidable opportunity to develop better, unified models that one day could cover the whole spectrum of tropical forests, both primary and managed.

## Author contributions

Conceptualization: E.D.; Methodology: E.D.; Software: A.D., E.D.; Experiments: A.D., E.D.; Formal analysis: E.D.; Writing: A.D., E.D.

## Acknowledgements

Authors would like to thank the anonymous referees for their comments that helped clarify the contributions and perspectives on the work. We also thank the editor for her professional handling of our manuscript and for bringing authors and referees on the same page to collectively produce scientific work of quality.

Illustrations in Figure 2 composed using free to use images by *macrovector* on Freepik.

## Appendix A Code and Models availability

The code used to create the experiments and generate the tables and figures reported in this paper is openly available at https://github.com/PUR-Projet/contextual_allometric_models (version 1.0.0), released under a CC BY-NC-SA 4.0 license. The Python environment is locked with the Poetry dependency manager, so the reported results can be reproduced exactly:

- `minimal_cofarm.py`reproduces the contextual modeling experiments
- `minimal_bound.py`reproduces the shifting context experiments

This open code policy allows also to retrain or fine tune proposed models on new data for further research.

The repository also provides frozen weights for all models in the `model_weights` directory and an easy to use script `prediction.py` that applies the pre-trained models to new data along with a minimal example. As such practitioners are empowered to apply these new models without the burden of training them on their own.

## Appendix B Additional Results

**Table B1.** Forecast of additional biomass prediction error due to covariate shift (test dataset). Arrows represent the shift direction: default →, reversed ←.

| Shift | $E[AGB_s]$ | $E[AGB_t]$ | $\Delta Z$ | $\mathcal{H}Disc$ | $\eta$ | $MAE_s^c$ | $MAE_t^c$ | $\Delta_{s,t} MAE$ | $\hat{\Delta}_{s,t} MAE$ |
| --- | --- | --- | --- | --- | --- | --- | --- | --- | --- |
| altitude-high⟵ | 661.9 | 857.6 | +3598.9 | 913.7 | 447.9 | 171.7 | 271.6 | +99.8 | +92.2 |
| altitude-high⟶ | 857.6 | 661.9 | -3598.9 | 1070.9 | 447.9 | 296.1 | 208.1 | -87.9 | +10.0 |
| altitude-low⟵ | 772.4 | 371.0 | -8793.6 | 3414.6 | 276.9 | 220.8 | 53.8 | -167.0 | -59.3 |
| altitude-low⟶ | 371.0 | 772.4 | +8793.6 | 2198.8 | 276.9 | 53.0 | 215.4 | +162.4 | +122.6 |
| continent-asia⟵ | 594.4 | 912.7 | +6569.2 | 3369.5 | 359.8 | 197.3 | 193.7 | -3.6 | +152.1 |
| continent-asia⟶ | 912.7 | 594.4 | -6569.2 | 1611.0 | 359.8 | 165.0 | 174.1 | +9.1 | -6.6 |
| drymonths-few⟵ | 621.0 | 1210.6 | +21406.9 | 8809.3 | 638.8 | 167.7 | 598.3 | +430.6 | +312.8 |
| drymonths-few⟶ | 1210.6 | 621.0 | -21406.9 | 7225.7 | 638.8 | 291.1 | 175.4 | -115.7 | -156.7 |
| drymonths-many⟵ | 700.1 | 182.8 | -11553.6 | 4104.4 | 245.4 | 186.2 | 56.7 | -129.6 | -81.2 |
| drymonths-many⟶ | 182.8 | 700.1 | +11553.6 | 12092.9 | 245.4 | 65.5 | 405.6 | +340.2 | +428.5 |
| foresttype-dry⟵ | 760.7 | 257.7 | -11134.2 | 4350.1 | 273.7 | 200.7 | 67.5 | -133.2 | -85.0 |
| foresttype-dry⟶ | 257.7 | 760.7 | +11134.2 | 1718.7 | 273.7 | 67.8 | 199.1 | +131.3 | +111.0 |
| foresttype-mangrove⟵ | 704.5 | 222.7 | -10262.3 | 3695.8 | 226.9 | 191.6 | 35.6 | -156.0 | -67.1 |
| foresttype-mangrove⟶ | 222.7 | 704.5 | +10262.3 | 6656.6 | 226.9 | 33.6 | 207.4 | +173.9 | +244.6 |
| foresttype-moist⟵ | 573.3 | 737.8 | +313.1 | 92.6 | 363.2 | 136.9 | 199.6 | +62.7 | +73.9 |
| foresttype-moist⟶ | 737.8 | 573.3 | -313.1 | 489.1 | 363.2 | 207.6 | 216.3 | +8.7 | +28.3 |
| foresttype-wet⟵ | 615.5 | 1064.0 | +16042.0 | 7068.6 | 506.7 | 172.5 | 458.0 | +285.5 | +257.2 |
| foresttype-wet⟶ | 1064.0 | 615.5 | -16042.0 | 3090.1 | 506.7 | 235.8 | 185.4 | -50.4 | -50.1 |
| oldgrowth-0⟵ | 763.5 | 234.5 | -11685.8 | 4332.3 | 247.2 | 208.3 | 35.9 | -172.4 | -84.5 |
| oldgrowth-0⟶ | 234.5 | 763.5 | +11685.8 | 4628.4 | 247.2 | 43.2 | 250.0 | +206.8 | +185.9 |
| rainfall-high⟵ | 621.5 | 986.1 | +13746.6 | 6027.8 | 477.7 | 175.6 | 421.6 | +246.0 | +225.8 |
| rainfall-high⟶ | 986.1 | 621.5 | -13746.6 | 2824.7 | 477.7 | 217.1 | 186.7 | -30.5 | -42.5 |
| rainfall-low⟵ | 788.9 | 322.2 | -11514.4 | 4312.0 | 308.9 | 199.9 | 100.5 | -99.3 | -83.9 |
| rainfall-low⟶ | 322.2 | 788.9 | +11514.4 | 3965.9 | 308.9 | 101.0 | 242.9 | +141.9 | +167.9 |
| random-1⟵ | 538.3 | 588.3 | -99.3 | 106.8 | 341.6 | 183.0 | 159.5 | -23.5 | +40.6 |
| random-1⟶ | 588.3 | 538.3 | +99.3 | 126.8 | 341.6 | 158.3 | 183.0 | +24.7 | +74.6 |
| random-2⟵ | 676.3 | 754.5 | +3138.5 | 1399.9 | 386.3 | 175.8 | 214.0 | +38.2 | +103.4 |
| random-2⟶ | 754.5 | 676.3 | -3138.5 | 1488.6 | 386.3 | 210.4 | 173.8 | -36.6 | -2.8 |

